# Utility and validity of group atlas versus personalized functional network approaches for depressive constructs

**DOI:** 10.64898/2026.03.10.710919

**Authors:** Ellyn R. Butler, Lauren B. Alloy, Damon D. Pham, Noelle I. Samia, Robin Nusslock, Amanda F. Mejia

**Affiliations:** Department of Psychology, Northwestern University, Evanston, IL 60208; Department of Psychology and Neuroscience, Temple University, Philadelphia, PA 19122; Department of Statistics, Indiana University, Bloomington, IN 47405; Department of Statistics and Data Science, Northwestern University, Evanston, IL 60208; NSF-Simons AI Institute for the Sky (SkAI), Chicago, IL 60611; Institute for Policy Research, Northwestern University, Evanston, IL 60208

**Keywords:** group, personalized, connectivity, fmri, validity, depression

## Abstract

**Background:** To understand the neurobiology underlying psychopathology, we need valid measurements of brain function. Group atlases for brain functional connectivity (FC) allow for efficient comparisons, but they fail to account for inter-individual variability in spatial features of networks, a problem that personalized approaches address. We assess the validity and predictive utility of group and personalized approaches of quantifying FC by 1) comparing effect sizes of associations with psychological metrics; and 2) accounting for spatial features of brain networks when examining the association between FC and psychological metrics.

**Methods:** 324 teens ages 13-16 participated. The personalized approach for estimating networks was a hierarchical Bayesian model. Effect size comparisons were done by comparing the correlations between FC estimated using different methods (group, personalized, and the intersection of the group and personalized maps) and psychological metrics (depression, ruminative coping style, and sensitivity to punishment/reward) with Steigler’s Z-test. We also conducted regressions, with psychological metrics as the dependent variables. Those models included FC and spatial features, together and alone.

**Results:** The effect size comparisons did not survive FDR correction. However, exploratory permutation tests show that 1) the magnitude of the correlations with depression are larger on average for the intersection estimates of FC than the group estimates; and 2) the magnitude of the correlations with a ruminative coping style are larger on average for the intersection estimates of FC than the personalized estimate. The other comparisons conducted using permutation tests are not significant. Multiple regression analyses demonstrated that only spatial features of networks, not FC, are associated with sensitivity to reward.

**Discussion:** These results imply that the intersection estimates are more valid than the group estimates, and that the intersection estimates have greater predictive utility than personalized estimates. Further, spatial features of networks may be useful in and of themselves in certain contexts. Therefore, researchers in psychiatry should take into consideration spatial features of networks to gain a better understanding of the neurobiology underlying psychopathology.

## 1 Introduction

Understanding the pathophysiology underlying psychopathology is a central goal of researchers and physicians alike [1]. In order to do this well, we must have valid measurements of key biological features [2], including aspects of brain function. Measures are valid when they tap into the construct as it has been defined. For instance, if we define depression as low mood or loss of interest or pleasure [3], then a valid measure would ask about aspects of these symptoms, as opposed to, for instance, one’s intelligence.

In the field of cognitive and affective neuroscience, the focus historically has been on the reliability of different metrics of brain function, as opposed to the validity of the metrics. This hinders our ability to understand the pathophysiology and biological mechanisms underlying diseases and disorders [4]. Therefore, we must work towards establishing and ultimately improving the validity of brain metrics that show promise in helping us better understand pathophysiology and mechanisms in order to eventually impact health outcomes.

Functional connectivity (FC) derived from functional magnetic resonance imaging (fMRI) data has been used to study human brain networks for over 30 years [5], and is increasingly investigated in the context of human health and disease [6, 7, 8, 9, 10, 11]. Recognizing that there are many common features in functional brain networks (hereafter referred to as “networks”) across people, researchers have developed group atlases, which group locations (e.g., vertices, voxels) in the brain into a fixed number of networks [12, 13, 14]. These group atlases then can be used by researchers to summarize high-dimensional functional data, improving our ability to compare results across studies, and to build theories about the nature of different networks.

Although there are common features in brain networks across people, there also is substantial inter-individual variation in the placement, size, and shape of networks – the spatial features of networks [15]. This suggests that analyses using group atlases mix in signal from different brain networks into any given network [16]. For instance, signal from the dorsal attention network may end up contaminating the estimates of visual network FC, as these networks are adjacent to each other in many group atlases [12, 13]. Therefore, results finding differences in FC across groups or along continuous dimensions using group atlases may observe such results due to differences in spatial features networks across groups and dimensions, rather than true differences in FC [16, 17].

This problem has led to the development of personalized functional network approaches, which classify locations in the brain as members of certain networks for each individual person. Using these methods, researchers have shown that functional brain networks are dominated by stable group and individual factors [15, 18], with spatial features of networks being largely stable across time and cognitive states [15]. Further, the proportion of the cortical surface devoted to a particular network – or the *expansion* of the network (which can only be estimated using personalized approaches as group atlases assume that networks have identical spatial features across people) – has been shown to be associated with related cognitive functions. For instance, the salience network has been shown to be more expansive among those who are more depressed [19], and greater fronto-parietal expansion has been shown to be associated with better cognitive function [20]. Therefore, it is important that the field increasingly move towards studying functional brain networks using personalized approaches.

Quantifying aspects of networks using a personalized approach is complicated by the fact that this is an emerging field, and as such, there are a variety of methods that differ substantially, with no single gold standard. One key consideration when choosing a method is whether or not to assume that each location in the brain belongs to exactly one network, or if each point should be allowed to belong to any number of networks. Previous work demonstrates that there are regions in the brain known as hubs that are critical for communication between networks, and as such, can be considered to belong to multiple networks [21, 22, 18]. However, popular methods such as Infomap and the multi-session hierarchical Bayesian model (MS-HBM) constrain points to belong to exactly one network [23, 15, 24, 25]. To address this problem, researchers have developed personalized functional network methods that allow points in the brain to contribute to any number of networks [26, 27, 28, 18, 29]. The network maps that emerge provide evidence in favor of relaxing the assumption that locations in the brain must belong to exactly one network. Using methods that allow for networks to overlap, researchers have found that there are in fact hubs at the individual level [18]. To provide the most valid representation of spatial features of networks, it is important to allow for each location in any given individual’s brain to belong to any number of networks.

Nevertheless, most prior studies examining the relationship between cognitive or psychological metrics and FC are based on methods that constrain each point in the brain to belong to a single network. These studies have shown mixed results on the predictive utility of and effects sizes associated with FC estimated using personalized functional network approaches (hereafter referred to as “personalized approaches”) in comparison to group atlas-based approaches (hereafter referred to as “group approaches”) (see Table 1) [25, 30]. Some have found larger effects for FC estimates based on personalized approaches [25], whereas others have found larger effects using group approaches [30]. The larger effects sometimes found using group approaches have at least two possible explanations: effects of mixing in FC from other networks, or of noise introduced by personalized approaches. For instance, considering people who are depressed have more expansive salience networks than those who are not [19], if the Yeo17 group atlas [12] was used to estimate FC, signals from adjacent networks such as the control networks actually could be originating from the salience network among depressed individuals. Therefore, any associations between depression and control network FC may be driven by group differences in spatial features of the salience network. This signal mixing effect may explain why bigger effects are sometimes found with group atlas approaches to estimating FC. Alternatively, the group atlas approaches simply may have less noise in the spatial features underlying FC estimates, as group atlases typically are an amalgamation of hundreds or thousands of participants’ data, whereas network membership derived from personalized approaches relies heavily on a single participant’s data.

**Table 1:** Glossary of key terms.

| Term | Definition |
| --- | --- |
| Functional connectivity (FC) | The correlation in the blood-oxygen level-dependent time series within and between networks. See Equations (1) and (2) for details. |
| Utility | The extent to which a metric of interest (e.g., group estimates of FC) is associated with other variables of interest. |
| Validity | The extent to which a metric of interest is measuring the construct of interest. |
| Personalized (“functional network”) approach | An approach to quantifying FC within and between networks that allows for the spatial features of networks to vary across people. |
| Group (“atlas-based”) approach | An approach to quantifying FC within and between networks that does not allow for the spatial features of networks to vary across people. |
| Intersection approach | An approach to quantifying FC within and between networks that only considers the vertices that are deemed members of a network by a personalized approach and are considered part of that same network by the group atlas. |

**Table 2:** Sample characteristics by sex.

|  | Female<br>(N=192) | Male<br>(N=132) | Overall<br>(N=324) |
| --- | --- | --- | --- |
| **Age** |  |  |  |
| Mean (SD) | 15.5 (1.04) | 15.4 (1.09) | 15.5 (1.06) |
| Median [Min, Max] | 15.5 [13.0, 18.1] | 15.4 [13.1, 17.5] | 15.5 [13.0, 18.1] |
| **Race** |  |  |  |
| African American or Black | 57 (29.7%) | 22 (16.7%) | 79 (24.4%) |
| Asian-American or Asian | 7 (3.6%) | 7 (5.3%) | 14 (4.3%) |
| Biracial/Multiracial | 10 (5.2%) | 13 (9.8%) | 23 (7.1%) |
| Caucasian/White | 116 (60.4%) | 89 (67.4%) | 205 (63.3%) |
| Native Hawaiian or Other Pacific Islander | 1 (0.5%) | 0 (0%) | 1 (0.3%) |
| Other | 1 (0.5%) | 1 (0.8%) | 2 (0.6%) |
| **Ethnicity** |  |  |  |
| Hispanic or Latino/a | 13 (6.8%) | 19 (14.4%) | 32 (9.9%) |
| Not Hispanic or Latino/a | 177 (92.2%) | 111 (84.1%) | 288 (88.9%) |
| Missing | 2 (1.0%) | 2 (1.5%) | 4 (1.2%) |
| **Average Caregiver Years of Education** |  |  |  |
| Mean (SD) | 16.5 (2.32) | 17.1 (2.45) | 16.7 (2.39) |
| Median [Min, Max] | 17.0 [3.00, 22.5] | 17.0 [12.0, 24.0] | 17.0 [3.00, 24.0] |
| Missing | 7 (3.6%) | 6 (4.5%) | 13 (4.0%) |
| **Family's Gross Total Income** |  |  |  |
| \$0 - \$4,999 | 1 (0.5%) | 0 (0%) | 1 (0.3%) |
| \$5,000 - \$19,999 | 2 (1.0%) | 0 (0%) | 2 (0.6%) |
| \$20,000 - \$34,999 | 6 (3.1%) | 4 (3.0%) | 10 (3.1%) |
| \$35,000 - \$49,999 | 11 (5.7%) | 3 (2.3%) | 14 (4.3%) |
| \$50,000 - \$74,999 | 22 (11.5%) | 7 (5.3%) | 29 (9.0%) |
| \$75,000 - \$99,999 | 18 (9.4%) | 8 (6.1%) | 26 (8.0%) |
| \$100,000 - \$149,999 | 25 (13.0%) | 25 (18.9%) | 50 (15.4%) |
| \$150,000 - \$199,999 | 28 (14.6%) | 24 (18.2%) | 52 (16.0%) |
| \$200,000 - \$249,999 | 22 (11.5%) | 15 (11.4%) | 37 (11.4%) |
| \$250,000 - \$299,999 | 16 (8.3%) | 15 (11.4%) | 31 (9.6%) |
| \$300,000 and higher | 25 (13.0%) | 15 (11.4%) | 40 (12.3%) |
| Not known | 2 (1.0%) | 2 (1.5%) | 4 (1.2%) |
| Missing | 14 (7.3%) | 14 (10.6%) | 28 (8.6%) |

Note: 12 years of education corresponds to having completed high school. Average Caregiver Years of Education is the average across two caregivers if the child has two caregivers, and is the years of education of a single caregiver if they have one caregiver.

**Table 3:**
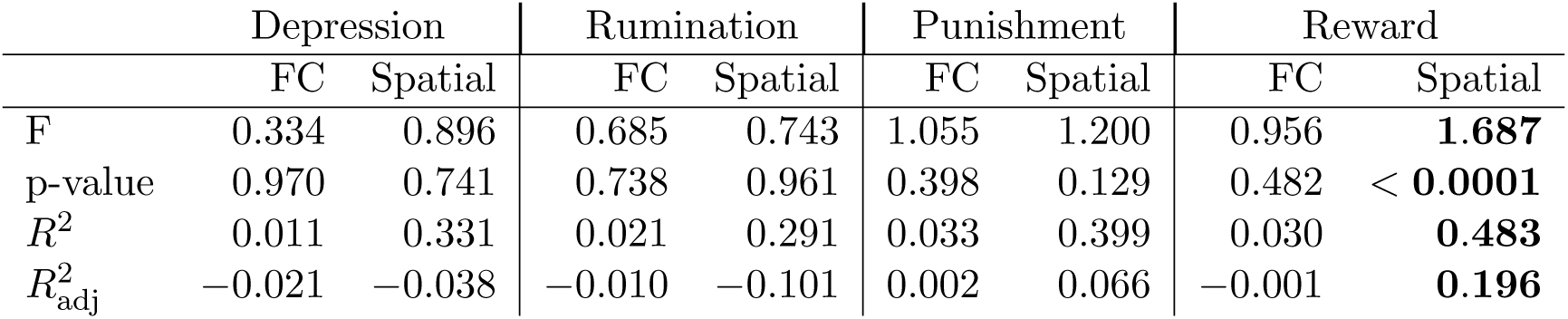
Full model results for group estimates of functional connectivity (FC) and spatial predictors, for each of the four clinical measures we consider (depression, ruminative coping style, sensitivity to punishment, and sensitivity to reward). To test each collection of predictors, we perform an F-test on all the corresponding coefficients. The first row shows the value of the *F* (10, 313) test statistic, and the second row shows the corresponding p-value.

It is important to keep in mind that the value of the greater validity provided by personalized approaches depends on the research goals. If the goal of the researcher is simply to predict some phenotype, a group atlas approach to estimating FC could be preferable to a personalized network approach if it has greater predictive utility. But if the goal is to *understand* something about the network, a personalized approach likely is preferable, as it does not mix signals across networks, and consequently, we are able to interpret FC estimates as originating from the selected network. Personalized approaches also can be expected to produce larger effect sizes in cases where there are true group differences in FC within a given network, barring excessive noise, given that they do not mix signals between networks. In other words, FC estimated using personalized approaches is more valid than FC estimated using group approaches, and as such, provides more interpretable findings.

In the current work, we evaluate whether signal mixing due to individual differences in spatial features of networks drives the higher predictive utility sometimes found with group approaches to estimating FC. To do so, we estimate FC within and between networks using three different network-defining approaches (see Figure 1 and Table 1): 1) the canonical Yeo17 group atlas [12]; 2) a personalized approach to estimating networks mapped onto the Yeo17 networks [27]; and 3) the *intersection* of each individual’s networks estimated using a personalized approach (hereafter referred to as “personalized networks”) and the group atlas. The intersection approach allows one to mask out locations in the brain that belong to other networks according to the group atlas, and potential noise from the individual networks, while excluding locations that are not part of a specific network for a given individual. Therefore, the intersection approach only retains features specific to the individual, while benefiting from the greater reliability of group atlases.

**Figure 1:**
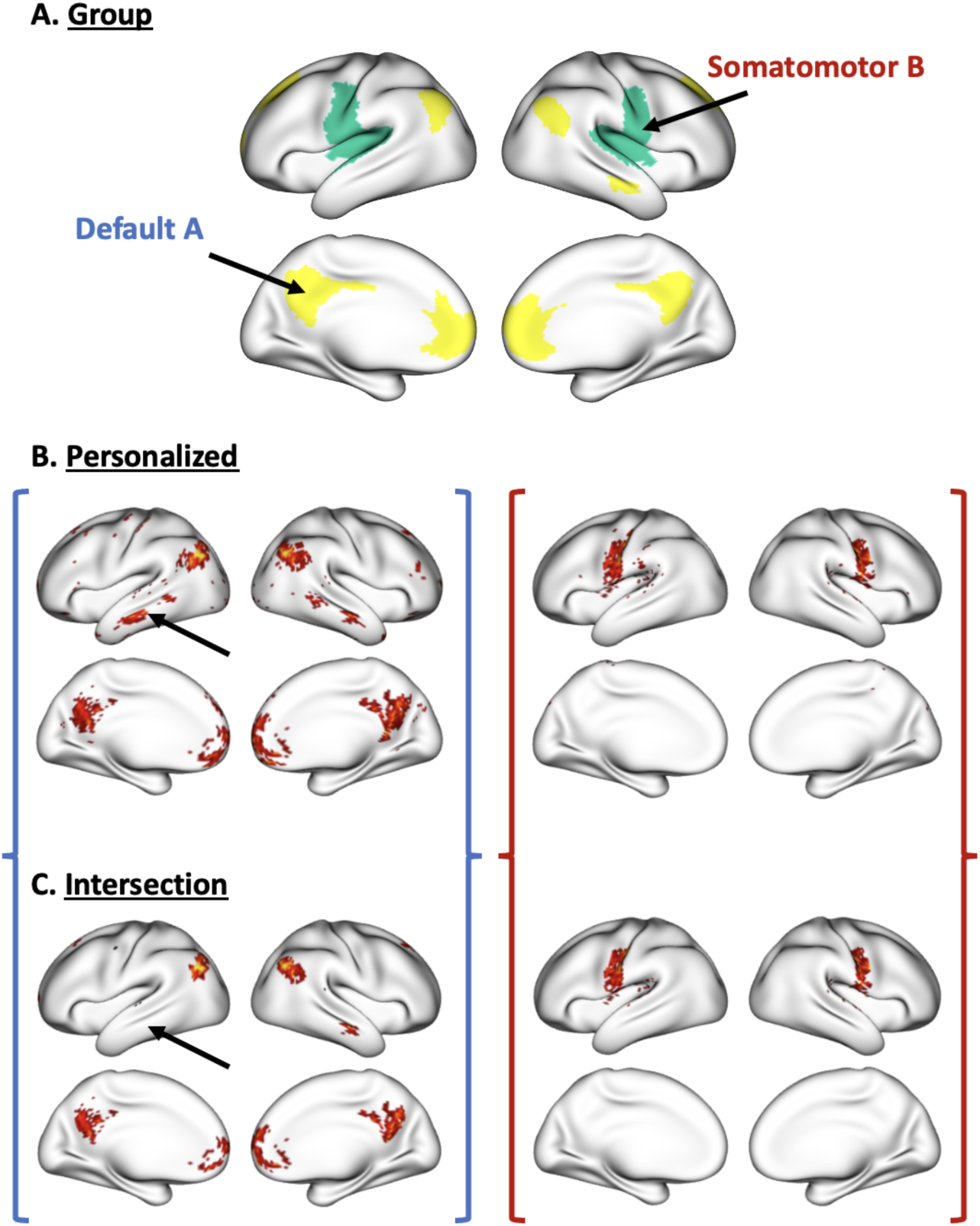
Group, personalized, and intersection maps. (A) shows the Default A and Somatomotor B networks isolated from the Yeo17 group atlas; (B) shows personalized thresholded engagement maps for an individual; and (C) shows intersection maps for this same individual. Black arrows in (B) and (C) point out differences between the personalized and intersection thresholded engagement maps.

By estimating FC using the intersection between the group atlas and each individual’s personalized networks, we are able to determine if the greater predictive utility observed with FC estimates using a group atlas is due to the mixing in of FC from other networks due to individual differences in the spatial features of networks. Specifically, we test the following hypotheses: Hypothesis #1 – if effect sizes with psychological metrics are greater for group atlas-based estimates of FC (hereafter referred to as “group estimates”) than intersection estimates, then what is useful about the group estimates is that they are mixing in FC across networks within a single network FC estimate, considering the primary difference between intersection and group estimates is that intersection estimates are based on maps that exclude vertices that belong only to the group atlas (i.e., are not classified as part of a given network by the personalized approach). Therefore, group estimates would have higher predictive utility and low validity. Hypothesis #2 – if effect sizes with psychological metrics are greater for the intersection estimates than the personalized estimates, then the personalized estimates have excessive noise. Therefore, personalized estimates of FC would have higher validity and low predictive utility. To provide greater clarity on the similarity in results across methods of quantifying FC, exploratory analyses are conducted to compare the effect sizes between the group and personalized estimates of FC.

Finally, we examine the extent to which FC estimated using a group atlas is associated with psychological metrics above and beyond spatial features. This directly tests whether the group atlas effects typically conducted in the literature are truly FC effects, or if they are driven by spatial features of networks. We use the following psychological metrics across analyses: depression, ruminative coping style, sensitivity to punishment, and sensitivity to reward. These were selected to maximize power in the current study, as these measures are available for the most participants. Although we focus on psychological metrics in the current work, these questions are relevant to brain-behavior associations using other phenotypes.

## 2 Methods

### 2.1 Participants

For this project, we use the Reward and Immune Systems in Emotion (RISE) [31] and Circadian, Reward, and Emotion Systems in Teens (CREST) datasets [32]. These datasets include fMRI and self-report data, in addition to a broad array of other metrics that are not the focus of this paper. Since July 2021, adolescents have been being recruited from the Philadelphia area to participate in the studies. To be eligible for RISE, participants must be 13-16 years old and not have experienced a major depressive episode. Further, participants are excluded from RISE if they have a history of cancer, heart disease, surgery, an ongoing autoimmune disease or disorder involving chronic inflammation (e.g., Crohn’s Disease, Type 1 Diabetes, Lupus, asthma, severe allergies), if they are HIV positive, or if they took immunosuppressant medications in the past three months.

To be eligible for CREST, participants must be 13-16 years old and not have experienced a manic or hypomanic episode or a bipolar spectrum disorder. Further, participants are excluded from CREST if they endorse any of the following: 1) use of melatonin or photosensitizing medications, 2) current or past treatment with bright light, or 3) shift work or travel across multiple time zones within the past month. These exclusion criteria are assessed in a phone interview and are designed to ensure that there are no contraindications or major confounding factors for the dim light melatonin onset (DLMO) procedures conducted in CREST. If a participant has traveled across time zones, but is otherwise eligible, participation is delayed for at least one month. Participants are excluded from both RISE and CREST if they endorse current psychotic symptoms, or have MRI contraindications (e.g., metal in the body, traumatic brain injury, pregnancy, severe claustrophobia; assessed with a standard MRI screening form). Participants in RISE are oversampled from the low end of the reward responsiveness dimension, whereas participants in CREST are oversampled from the high end [33, 34]. All participants and caregivers provided informed consent/assent. Temple University’s Institutional Review Board approved both protocols.

The final sample consists of adolescents who have complete depression, ruminative coping style, sensitivity to punishment, sensitivity to reward, and neuroimaging data as of June 18, 2025. Neuroimaging data are considered complete for a given participant if they have a T1-weighted structural scan, at least five minutes worth of degrees of freedom remaining following confound regression from the combined functional scans, and have a mean framewise displacement less than 0.5mm. 473 adolescents at least began the screening process (i.e., answered at least the first question from the Sensitivity to Punishment and Sensitivity to Reward Questionnaire (SPSRQ)). 82 adolescents were missing some data from at least one of the clinical measures (40 missing depression; 32 missing ruminative coping style; and 34 missing sensitivity to punishment or reward). Four adolescents had some neuroimaging data but had not started the SPSRQ (i.e., are not part of the 473). 62 adolescents did not participate in the neuroimaging session. No adolescents had fewer than five minutes of data remaining after confound regression (range = 8.58 - 29.04 minutes; mean = 27.53 minutes; median = 28.55 minutes). 29 adolescents had a mean framewise displacement greater than 0.5mm. 149 adolescents did not have sufficient clinical or neuroimaging data, resulting in a final sample of 324 adolescents.

### 2.2 Assessment

#### Depression

We use the Beck Depression Inventory - II (BDI-II) to assess depression severity over the preceding two weeks [35]. The BDI-II taps into constructs of sadness, pessimism, suicidal thoughts or wishes, loss of interest, worthlessness, and difficulty with concentration, among other depression-related constructs. Historically, the BDI-II has been shown to have a two-factor structure [36, 37] – comprising factors tapping into cognitive-affective and somatic-vegetative symptoms of depression. Exploratory factor analyses on the current sample demonstrate a largely unidimensional structure (λ_1_ = 4.65, λ_2_ = 1.15, *ε* = 0.86, *ω_H_* = 0.69), which implies good internal consistency and justifies using a sum score of the items [38]. The BDI-II has demonstrated good psychometic properties, including high test-retest reliability (*r* = 0.73 to 0.96), and convergent validity (i.e., high correlations with other measures of depression) [37].

#### Ruminative coping style

The 10-item Ruminative Responses Scale (RRS) is used to assess tendency to use a ruminative coping style [39]. The RRS taps into constructs of thinking about how alone one feels, how fatigued one is, how one wishes a situation had gone better, and how angry one is with oneself, among other rumination-related constructs. Historically, the RRS has been shown to have a two-factor structure [39] – reflective pondering and brooding. Exploratory factor analyses on the current sample demonstrate a largely unidimensional structure (λ_1_ = 8.48, λ_2_ = 1.27, α = 0.92, *ω_H_* = 0.75), which implies good internal consistency and justifies using a sum score of the items [38]. The RRS has demonstrated decent psychometric properties, including moderate test-retest reliability (*r* = 0.60) [39].

#### Sensitivity

We use the Sensitivity to Punishment and Sensitivity to Reward Questionnaire (SPSRQ) to assess sensitivity to punishments and rewards [34]. The SPSRQ taps into constructs such as refraining from doing illegal activities, being motivated by money, doing things to be praised, and showing off in a group. Historically, the SPSRQ has been shown to have a two-factor structure [34] – sensitivity to punishment and sensitivity to reward. Exploratory factor analyses on the current sample demonstrate a largely two-dimensional structure (λ_1_ = 6.36, λ_2_ = 4.56, α = 0.83, *ω_H_* = 0.52), which justifies using separate sum score of the items for sensitivity to punishment and sensitivity to reward, respectively [38]. The SPSRQ has demonstrated good psychometric properties, with high test-retest reliability at three months for the punishment (*r* = 0.89) and reward (*r* = 0.87) subscales [34].

### 2.3 Neuroimaging

#### Acquisition

Imaging data were collected at the Temple University Brain Research & Imaging Center (TUBRIC) on a Siemens MAGNETOM Prisma 3-Tesla scanner with a 64-channel phased-array parallel transmit and receive RF head/neck coil. Structural images were acquired using an MPRAGE sequence to acquire 208 axial slices (voxel size = 0.8mm^3^; TR=2300ms; TE=2.99ms; FOV=256×256; Matrix=320×320; Flip Angle=7°). Three functional scans were used: The Chatroom Interact task [40], and two runs of the Monetary Incentive Delay (MID) task [41]. The Chatroom Interact task uses a slice-accelerated multiband EPI sequence (multiband acceleration factor: 2. GRAPPA acceleration factor: 2) covering 64 axial slices (voxel size = 2.0mm^3^; TR=2050ms; TE=25ms; FOV=208×208mm; Matrix=104×104; Flip Angle=76°). The MID task also uses a slice-accelerated multiband EPI sequence with identical scan parameters. A total of 390 volumes were collected for the Chatroom Interact task, and 289 for each of the runs of MID.

These tasks can be concatenated to increase reliability of functional network metrics [42]. Further, functional network metrics can be validly and reliably estimated using task data [43, 44], and functional network metrics derived from tasks tend to have greater predictive utility than rest even when the task is not aligned with the phenotype of interest [43]. We therefore concatenate these three functional scans after processing for use in all analyses.

#### Processing

We use fMRIPrep version 23.2.0 [45], Connectome Workbench [46], and R packages [47, 48] to process the magnetic resonance imaging data. Processing steps include correcting T1-weighted images for intensity non-uniformity and skull-stripping; slice-time correcting the functional images, resampling them into their native space, and then projecting them into the fsLR-32k surface space (resampled to 10k per hemisphere for computational purposes). We use simultaneous nuisance regression (translation (x, y, z), rotation (yaw, pitch, roll), and global signal; the derivatives, squares, and squared derivatives of each of those signals; outlier flagging with DVARS (dual cutoff method) [49]; and .01 Hz high-pass filter using 12 discrete cosine transform bases) [50]. DVARS is estimated on otherwise nuisance-regressed data to reduce redundancy in the covariates [48].

#### Network metrics

To construct individual-level network engagement maps, which are then used to derive the personalized estimates of FC, we use a Bayesian Brain Mapping (BBM) technique [51] (Figure 2). BBM employs a hierarchical Bayesian framework with group-derived priors on the spatial features of networks. The role of the priors is to mitigate noise and produce reliable maps of individual vertex-level network engagement and membership [27]. In the current work, the priors are derived from the data used in the downstream analyses, resulting in an empirical Bayesian approach. Since the priors are established before model fitting, the BBM model can be fit for each individual separately for efficient computation. We use the Yeo17 atlas [12] as a template instead of group independent component analysis (ICA) maps as in previous work [27], to allow comparisons to FC estimates derived using the Yeo17 group atlas. The Yeo17 atlas was used as a template to guide the prior, such that it was assumed that there are 17 networks, which should resemble the topographical layout of the Yeo17 atlas, while being allowed to overlap.

**Figure 2:**
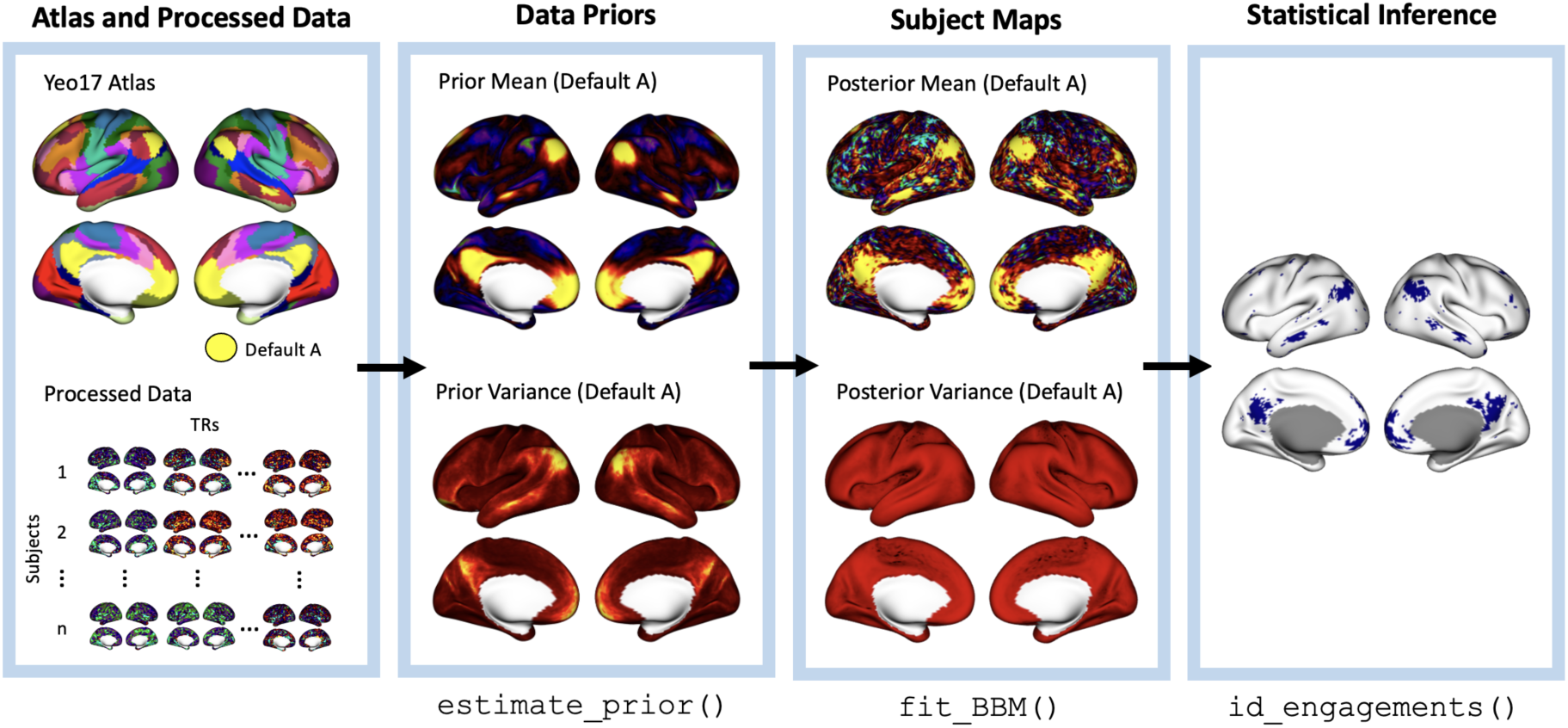
Pipeline for estimating personalized networks using the BayesBrainMap R package [51]. Functions that produce the surface outputs in the blue boxes are listed below their corresponding boxes. The Default A network is highlighted for illustration purposes. Features from all Yeo17 networks were estimated.

We derive prior mean and variance maps from the Yeo17 group atlas in combination with the processed functional data from each participant in the sample [51], similarly to how we have done previously [52]. Briefly, each functional scan is split down the middle to produce pseudo test-retest data, and for each pseudo-session we produce a map of network engagement for each of the 17 networks using a variation of dual regression [53]. In the first “regression” we compute the median time course across the vertices within each parcel, and in the second regression we regress those time courses against the fMRI time series to obtain maps of network engagement. Using the resulting set of test-retest network engagement maps for all participants, we estimate the prior mean and between-participant variance maps using variance decomposition [27]. We then use these prior mean and variance maps as priors in the BBM model for individual network maps.

After fitting the BBM model, we identify vertices belonging to each of the 17 networks by performing the Bayesian equivalent of a hypothesis test at every vertex, based on the estimated engagement level and its posterior standard deviation. Although in hypothesis testing it is common to test against a null hypothesis of zero engagement, here to avoid statistically significant but negligible effect sizes, we test against a null hypothesis of negligible-or-smaller engagement. For any vertices where the null hypothesis is rejected, we can conclude that they exceed a predetermined minimum effect size. Specifically, we set the minimum effect size to correspond to a z-score of *z* = 1 of the prior mean map. To avoid false positives, for each network we apply False Discovery Rate (FDR) correction [54] to control the FDR at 0.01 across all 18566 vertices.

After identifying vertices with significant engagement in each network, we compute inter- and intra-network FC. *Inter* -network FC is computed as the correlation between the time series for each network. The network time series are computed across all significant vertices in the network, weighted by the surface area that each vertex occupies in fsLR-32k space and the estimated engagement of the vertex for a given network. Specifically, the following equation was used to compute personalized FC (*C_ghi_*) between networks *g* and *h* = 1*, … ,* 17 for participant *i*:

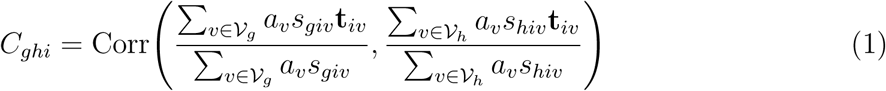

where *V_g_* indexes the vertices belonging to networks *g* = 1*, … ,* 17, and *h* = 1*, … ,* 17, respectively; *a_v_* = the surface area of the vertex *v* in fs-LR32k space; *s_giv_* = the engagement level for participant *i* in network *g* at vertex *v*; and **t***_iv_* = the time series for participant *i* at vertex *v*. The intersection estimates of inter-network FC are computed in a similar manner, except vertices that are not belonging to a given network in the Yeo17 atlas were excluded (Figure 1). *Intra*-network FC for each network *g*, *D_gi_*, is computed as the average correlation in the BOLD time series between all pairs of vertices in the network, weighted by the surface area that each vertex occupies in fsLR-32k space and the engagement of the vertex in network *g*:

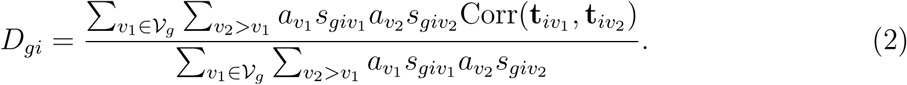

For group-based FC, the correlations between pairs of vertices only were weighted by the surface area of the vertices, as all vertices have equal engagement in the Yeo17 group atlas. Finally, we derive spatial and FC regressors for the multiple regression models. For the spatial covariates, we perform singular value decomposition on the engagement maps for each network separately. To reduce background noise, we first threshold the engagement maps such that negligible engagements are set to zero. Specifically, we apply a threshold of *z* = 2 on the prior mean map. Singular vectors that explain at least 1% of the variance in engagement were retained for the regressions. For the FC regressors, we compute intra- and inter-network FC using the Yeo17 atlas and perform singular value decomposition to reduce the dimensionality, and retain the singular vectors that explain at least 1% of the variance in FC.

### 2.4 Statistical Analyses

We conduct two sets of analyses to evaluate the utility and validity of group versus personalized approaches: 1) effect sizes comparisons of the association between intra- and inter-network FC estimated using group, personalized, and intersection network maps and a series of psychological metrics (effect size comparisons); and 2) ability of intra- and inter-network FC estimated using a group atlas to predict psychological metrics, above and beyond network spatial features of networks (multiple regressions).

#### Effect Size Comparisons

We perform comparisons of correlations between psychological metrics and FC estimated using a group atlas, personalized networks, and the intersection between the two. We use Steiger’s Z-test for equality of correlations [55], as it accounts for dependence between the observations across the two correlations. Specifically, for the *k*th clinical metric, *k* = 1*, … ,* 4, and *j*th FC estimate (inter- and intra-network FC), *j* = 1*, … ,* 153, *r*_1*jk*_ = the correlation between the metric and group-based FC; *r*_2*jk*_ = the correlation between the metric and personalized FC; and *r*_3*jk*_ = the correlation between the metric and intersection FC.

Dropping the *j* and *k* subscripts for simplicity momentarily, the test statistic *Z_AB_* for comparing correlations *r_A_* and *r_B_*, where *A* ≠ *B*, *A, B* = 1, 2, 3, takes the following form:

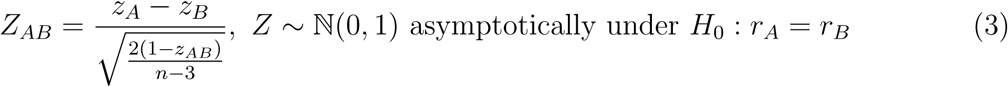

where *z_A_* and *z_B_*, are the Fisher-transformed correlations *r_A_* and *r_B_*, respectively, and *z_AB_* is the Fisher-transformed correlation between the two FC metrics of interest. To evaluate Hyp. #1, we test if *r*_1*jk*_ and *r*_2*jk*_ differ for each measure of inter- and intra-network FC *j* and metric *k*. That is, *r_A_*= *r*_1*jk*_, *r_B_*= *r*_2*jk*_, and *r_AB_* = *r*_12*j*_. To evaluate Hyp. #2 using the continuous self-report measures, we test if *r*_2*jk*_ and *r*_3*jk*_ differ for each network *j* and metric *k*. That is, *r_A_* = *r*_2*jk*_, *r_B_* = *r*_3*jk*_, and *r_AB_* = *r*_23*j*_. The null hypotheses are rejected based on the p-values from the Steiger test, controlling the False Discovery Rate at 0.01 across all 2142 tests to account for multiple comparisons [54].

#### Multiple Regressions

For each clinical metric *k* = 1*, … ,* 4, we fit a series of linear models using observations from *n* participants, where the dependent variable of interest is the clinical metric (*Y_ik_*), with error terms 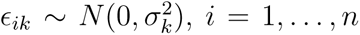. Independent variables include (a) the SVD singular vectors derived from the FC estimates based on the Yeo17 group atlas (***w****_i_*) and (b) the SVD singular vector scores from the spatial network engagement maps (***u****_ij_*). Recall that we perform a separate SVD on the engagement maps for each network, so ***u****_ij_* is indexed by both participants *i* and spatial singular vectors *j* = 1*, … , J*. We fit four separate sets of multiple regression models, one set for each clinical metric. For each metric, the full model takes the following form:

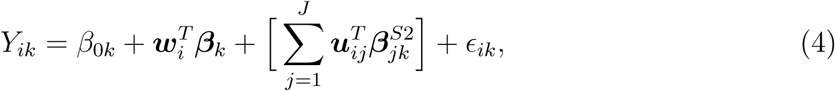

where *β_k_* is a vector of coefficients for ***w****_i_*, and 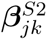 is a vector of coefficients for ***u****_ij_*.

The first reduced model includes only the spatial singular vectors and takes the following form. Comparing the full model to this reduced model allows us to determine if the group atlas-derived FC estimates (***u****_i_*) explain any additional variance in the psychological metrics above and beyond the spatial singular vectors (***u****_ij_*), and how much:

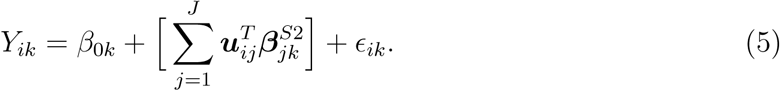

The second reduced model includes only the FC predictors and takes the following form. Comparing the full model to this reduced model allows us to determine if the spatial singular vectors explain a portion of the predictive power of the group atlas-derived FC estimates, and how much:

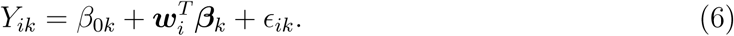

We use nested model *F* tests to compare the reduced and full models. We only perform these nested model tests for psychological metrics where the full model explained a significant amount of variance in the metric.

## 3 Results

In this section, we summarize the results of the effect size comparisons and the multiple regressions model comparisons to assess the validity and predictive utility of personalized and group-based approaches to quantifying FC. The effect size comparison analyses show that, after applying FDR correction, none of the individual effect size comparisons remain significant. However, a different picture emerges when examining the average difference in magnitude of the associations across all of the FC variables using permutation tests (see Supplement for details). Figure 3 shows that 1) the magnitude of the correlations with depression are larger on average for the intersection estimates of FC than the group estimates (*p* = 0.038); and 2) the magnitude of the correlations with a ruminative coping style are larger on average for the intersection estimates of FC than the personalized estimates (*p* = 0.010). These results imply that 1) the intersection estimates may remove signal from non-target networks that are included in the group estimates, which for the group estimates reduces observed effect sizes; and 2) the intersection estimates may remove noise that is often characteristic of personalized approaches. The other comparisons conducted using permutation tests are not significant.

**Figure 3:**
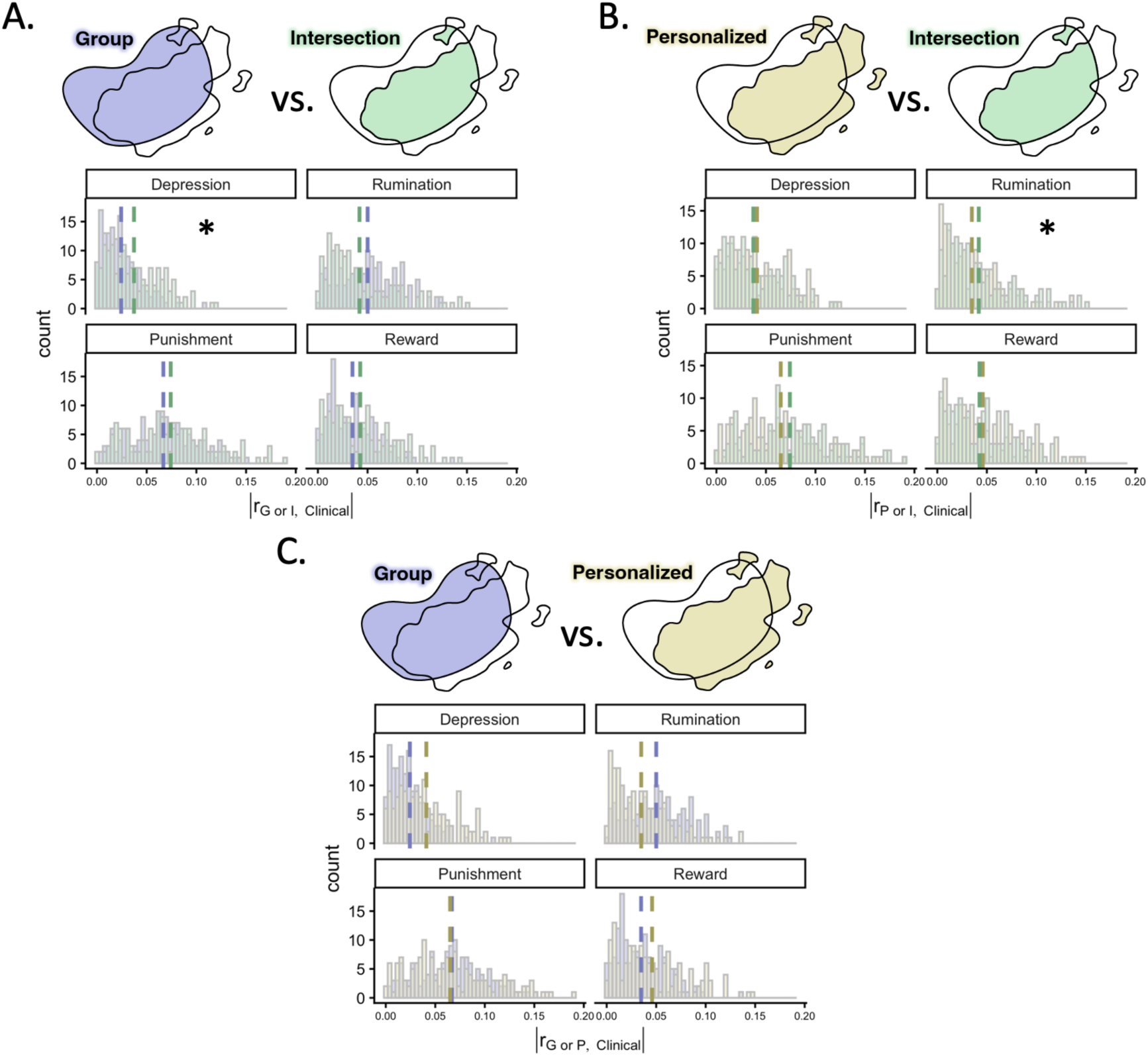
Histograms of the absolute values of the correlations between psychological metrics and different ways of quantifying FC. Comparisons are made to test if the absolute values of the correlations between FC quantified using one of the three methods (group, personalized, intersection) and the psychological metrics differed across methods. None of the comparisons survived FDR correction. Permutation tests then are used to determine if, on average, differences in the absolute values of the correlations differ on average across methods of quantifying FC for each clinical metric. Significant results are indicated with an asterisk. A.) Comparison of the correlations with psychological metrics for the group and intersection methods; B.) Comparison of the correlations with psychological metrics for the personalized and intersection methods; and C.) Comparison of the correlations with psychological metrics for the group and personalized approaches.

Turning now to the multiple regression models, 10 FC singular vectors (based on the group estimates of FC) and 115 spatial singular vectors are retained using the 1% variance explained cutoff for inclusion of singular vectors from the SVDs. The multiple regression analyses showed that neither group estimates of FC nor spatial features of networks explained a significant portion of variance in depression, ruminative coping style, or sensitivity to punishment, based on the full model. As such, we do not consider the nested models for these clinical measures.

Spatial features of networks, but not group estimates of FC, explain a significant portion of variance in sensitivity to reward, based on the reduced models. Using nested model comparisons, group estimates of FC did not explain additional variance in sensitivity to reward above and beyond spatial features (*F* (198, 208) = 0.84, *p* = 0.588). As a sensitivity analysis, we repeat the multiple regression models using a 2% variance explained cutoff for inclusion of singular vectors from the SVDs. Using this criterion, 5 singular vectors based on group estimates of FC and 28 spatial singular vectors are retained. Results of the full and reduced models using these features are consistent with those reported above.

## 4 Discussion

There is increasing evidence and support for the idea that accounting for the inter-individual variability in the spatial features of networks may result in more valid estimates of FC. Yet, both personalized and group approaches to estimating FC may have good predictive utility, depending on the outcome of interest. In the current study, we interrogate specific hypotheses around the advantages and disadvantages of group versus personalized approaches by comparing the predictive utility of three different methods for quantifying FC. In addition to group and personalized approaches to quantifying FC, we introduce a novel intersection-based approach designed to realize the advantages of both. Further, we interrogate whether the predictive utility of group estimates of FC is derived, at least in part, from mixing in signal from non-target networks due to inter-individual variability in spatial features of the networks.

After controlling for multiple comparisons over the 1836 comparisons of correlations between FC and psychological metrics, we find no statistically significant differences in the magnitude of the correlations between various ways of quantifying FC. However, after conducting a series of permutations tests, we see that 1) the magnitude of the correlations with depression are larger on average for the intersection estimates of FC than the group estimates; and 2) the magnitude of the correlations with a ruminative coping style are larger on average for the intersection estimates of FC than the personalized estimates. These results imply that 1) the intersection estimates may remove signal from non-target networks that are included in the group estimates, which for the group estimates reduces observed effect sizes, suggesting that the intersection estimates are more valid than the group estimates; and 2) the intersection estimates may remove noise that is often characteristic of personalized approaches, suggesting that intersection estimates have greater predictive utility than personalized estimates. The other comparisons conducted using permutation tests are not significant. These results suggest that, with respect to some of the clinical outcome metrics used in the current study, the intersection estimates may be the best in terms of both predictive utility and validity.

Using multiple regression models we find that, among the four psychological metrics tested, only spatial features explain a significant portion of the variance in sensitivity to reward, and for no other psychological metrics. The fact that spatial network features are associated with sensitivity to reward suggests that there are functional brain features that do meaningfully relate to clinical outcomes. Therefore, future studies may benefit from deriving spatial features of networks when using personalized approaches, instead of simply considering FC. This represents an opportunity unique to personalized approaches, as group atlases assume that spatial features of networks are invariant across people.

The current study is limited by a small set of outcome metrics and a relatively small number of participants, particularly given the high-dimensional nature of our functional network features. With only four clinical measures to serve as dependent variables, we are restricted in our ability to determine the relative predictive utility and validity of our FC metrics. Other potentially relevant dependent variables include performance on vocabulary, reading, and crystallized cognition tasks [56], and behaviors such as mobility as captured by smartphones [57]. The study also is limited by the fact that there are 324 participants in the current work, while we are trying to draw inferences about thousands of functional brain network features. Notably, this is a common problem in neuroimaging studies, with many studies having far fewer participants [58, 59], and we did conduct a variety of dimensionality reduction and multiple comparison correction procedures. Nevertheless, this remains a concern, as it is possible that with a more focused, theory-driven analysis we may have been able to find significant results.

Our study has several notable strengths. First, we propose a novel intersection approach to realize some of the advantages of both group and personalized approaches to estimating FC. The intersection approach allows us estimate FC only in locations (i.e., vertices) that belong to both the group and personalized maps. The intersection improves over the group atlas by excluding features that do not map onto the spatial features of networks for a given individual. It also may improve over the personalized network maps by capitalizing on the noise reduction properties of group atlases, although it runs the risk of missing important signal from hubs and true features of individuals’ networks. Therefore, our intersection approach may be useful for future studies examining the validity of methods for estimating FC. Further, intersection network maps may provide a reasonable compromise between noisy personalized maps and rigid group atlases for quantifying FC.

Finally, a strength of our study is our use of a Bayesian Brain Mapping (BBM) method that meaningfully improves over popular methods for estimating personalized networks [27]. Popular methods include dual regression [53] and Infomap [15, 42]. BBM improves over dual regression by reducing noise with population-derived priors on the spatial features of networks. BBM also improves over methods like Infomap by allowing vertices to be classified as belonging to multiple networks. This represents an important advantage of BBM, as we know that there are hubs – regions that affiliate with multiple networks – in the brain [60, 61, 22]. Notably, work that has compared personalized approaches that allow for network overlap to those that do not has found that allowing for network overlap produces more reliable maps than constraining locations in the brain to belong to a single network [29].

Future research examining the utility and validity of different methods of quantifying FC should include outcomes metrics that have been shown to be more robustly associated with FC. Generally speaking, FC is more strongly associated with cognitive metrics than psychological metrics [56], so it will be important to conduct similar analyses in a large dataset with participants who have cognitive measures. In such a dataset, we may gain greater clarity on the nature of what is useful about group and personalized estimates of FC. Our findings support the idea that future studies that seek to uncover the neurobiological bases of psychopathology should continue to investigate spatial features of networks, as these features are emerging as potentially important biological substrates of psychopathology [62, 19, 63, 52]. Therefore, whether or not personalized approaches provide more valid estimates of FC than group atlases, it will continue to be important to utilize personalized network approaches as they allow us to quantify spatial features of networks. More broadly, if we are able to validly measure these spatial features, we will be able to gain a better understanding of the neurobiology underlying psychopathology, opening the doors to identifying potentially effective targets for intervention.

## 5 Data and Code Availability

The final derived dataset can be found as part of the National Data Archives for RISE (C3835) and CREST (C4809), and code for this paper can be found in the accompanying GitHub repository: https://github.com/NU-ACNLab/personalized_versus_group/.

## 6 Author Contributions

E.R.B. conceptualized the work, conducted data analysis, and wrote the manuscript. A.F.M. supervised project conceptualization, execution, and data analysis. A.F.M. and D.D.P. assisted with image processing. N.I.S. provided feedback on data analysis. L.B.A. and R.N. obtained the funding for data collection, collected the data, and provided guidance on using the available data. All authors reviewed and edited the manuscript.

## 9. Acknowledgments

This research was supported in part through the computational resources and staff contributions provided for the Quest high performance computing facility at Northwestern University which is jointly supported by the Office of the Provost, the Office for Research, and North-western University Information Technology.

