## Supplement for "Utility and validity of group atlas versus personalized functional network approaches for depressive constructs"

### 10 Supplementary Materials

#### 10.1 Deviations from the Preregistration

- Neuroimaging exclusion criteria: We added an additional exclusion criteria for the neuroimaging data such that scans with mean framewise displacement greater than 0.5 mm were excluded. This was an oversight in the preregistration, as we did not appropriately anticipate having very high motion scans.
- Dimensions of depression and ruminative coping style: The Beck Depression Inventory - II and the 10-item Ruminative Responses Scale were both found to be unidimensional, and as such, single scores were derived from each measure. This deviates from the preregistration, as we expected to find a two-dimensional structure for each measure, as this has been found previously.
- Exclusion of the Pittsburgh Sleep Quality Index (PSQI): We decided to exclude the PSQI from analyses prior to any data analysis given its conceptual deviation from the other measures. Specifically, it does not directly tap into emotional or motivational constructs, unlike the other measures.
- Processing changes: Three changes were made to the processing stream to address noisy participant engagement maps: 1) lowering the $z$ threshold to 1; 2) concatenating additional functional data; and 3) calculating DVARs on residualized data. The effect size comparison analyses were initially run for the purposes of Flux Society's annual 2025 conference. In the process of producing results for this poster, we discovered that setting $z = 3$ as was initially proposed resulted in many participants having no vertices being determined to belong to specific networks. Upon visually inspecting some of the participants' engagement maps, we (Ellyn Butler and Amanda Mejia) determined that the estimates were far noisier than desired. To combat this, we decided to 1) lower the $z$ threshold to 1 to avoid overly restricted network engagement maps; 2) concatenate the Chatroom and Monetary Incentive Delay tasks to gain additional degrees of freedom;

and 3) calculate DVARs on the otherwise residualized functional data before conducting a single nuisance regression on the raw data to reduce redundancy in the motion artifact picked up by the DVARs-based flags and the motion regressors. These three changes resulted in all participants having at least some vertices deemed members of each network.

- Time series: Originally, we proposed partialing out the variance associated with other networks from the participants' time series ("Since the personalized networks can overlap, the network time series were computed using multiple linear regression in order to partial-out the contribution of other networks for the purposes of estimating inter-network FC."). We did not do this ultimately because we believe that weighting the vertex by its engagement in a specific network for a particular participant serves a similar purpose.
- Data-driven spatial features: We originally proposed to use principal components analysis on the engagement maps to derive spatial features to control for in multiple regression analyses, and then to use cross-validation to select components. We found a more computationally efficient method, however, which we now detail in the main text.
- Expansion as a covariate: We decided to exclude the network expansion estimates as covariates from the multiple regression models because these estimates were highly collinear with the first singular vectors from the singular value decompositions for each of the networks (Figure S4).

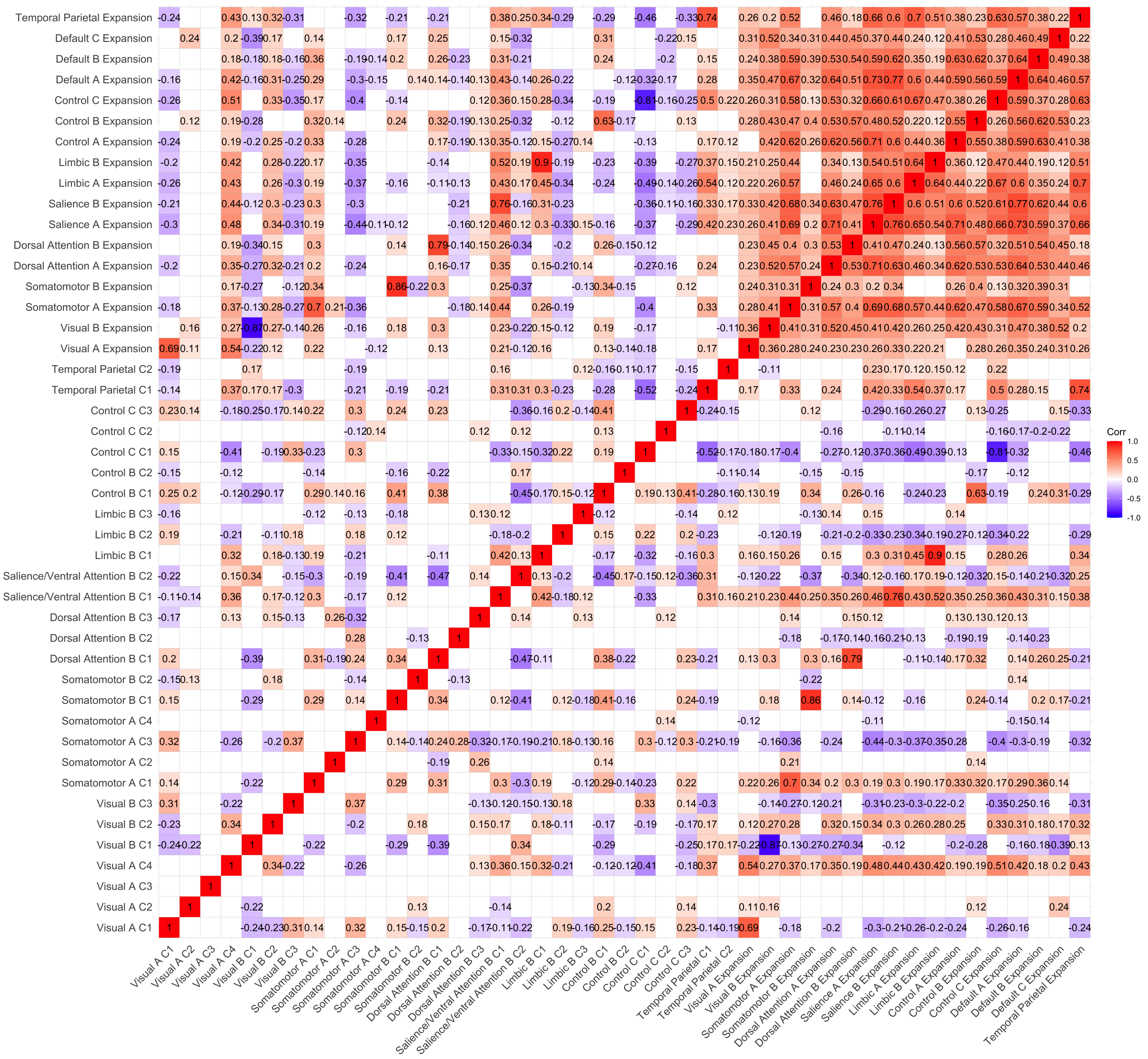

Figure 4: Correlations between expansion estimates and singular vectors explaining at least 2% of the variance. Depicted correlations are significant at the $\alpha = 0.05$ level. Singular vectors are delineated by “C”.

- Dimensionality reduction: In the preregistration, we stated that “we will perform a principal components analysis on each of the 17 network engagement maps across people, where the number of principal components to retain will be determined through cross validation. These principal components and network expansion will serve as control variables in the multiple regression analyses.” This statement is inherently vague, however, and as such, required further clarification. Upon inspecting the scree plots from singular value decompositions for both the engagement maps (Figure S5) and the FC estimates, we readily saw that only the first few singular vectors from each decomposition explained a notable amount of variance in the maps or FC values. Therefore, we used conservative thresholds of 1% and 2% variance explained for our regression analyses.

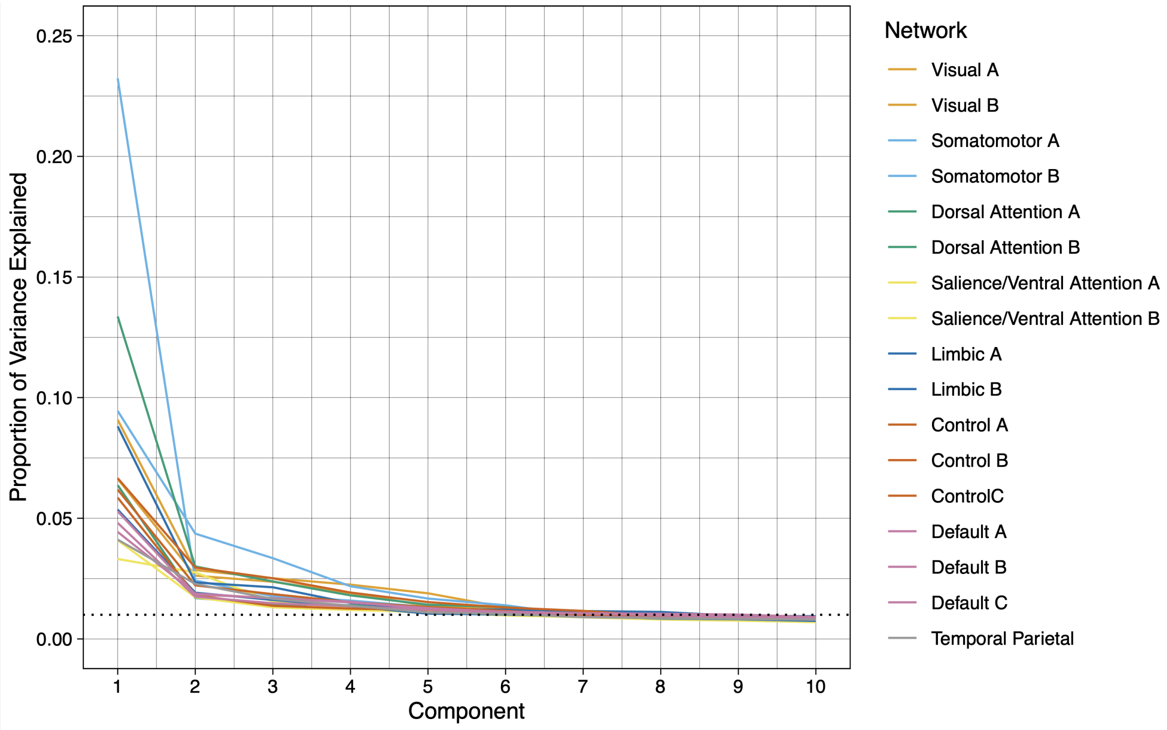


Figure 5: Proportion of variance explained in the engagement maps by the first ten singular vectors. The dotted line represents the cutoff of 1% for inclusion in the multiple regression models.

- Penalized regression: Given that we were able to reduce the dimensionality of the independent variables sufficiently by using a cutoff of 1% variance explained in the singular value decompositions, there was no need to implement a penalized regression. For sensitivity analyses, we ran the same analyses with a cutoff of 2% variance explained, which yielded analogous results.

#### 10.2 Permutation tests

For each pair of methods of quantifying FC (Group vs. Intersection, Personalized vs. Intersection, and Group vs. Personalized) and each clinical metric (depression, ruminative coping style, sensitivity to punishment, and sensitivity to reward), permutation tests are conducted. For each of 10,000 iterations, values for pairs of FC variables (e.g., Group Salience A - Salience B FC and Personalized Salience A - Salience B FC) were swapped for the same participants across all pairs of FC variables to maintain the dependency structure within participants. Swaps were made for participants if their row number corresponded to a randomly sampled number less than 0.5 from a uniform distribution from 0 to 1, resulting in, on average, half of the participants having their values swapped across permutations. For each permutation, the following statistic was calculated:

$$Q_{klms} = \frac{\sum_{j=1}^{153} |r_{jkl}| - |r_{jkm}|}{153}, \quad (7)$$

where $k = 1, \ldots, 4$ indexes the clinical metric; $l$ = the first method of quantifying FC; $m$ = the second method of quantifying FC; $s = 1, \ldots 10,000$ indexes the permutation; and $j = 1, \ldots, 153$ indexes the FC variable type (e.g., Salience A - Salience B FC). The $p$-value for each of these comparisons was estimated as the portion of permuted values that were more extreme than the observed value.

#### 10.3 Other analyses

- Ridge regressions were estimated using FC estimates from the Schaefer 400 atlas [14] using nested five-fold cross validation to choose the penalty parameter values and then obtain out of sample $R^2$ estimates. The decision to do this was made after the preregistration was posted and other analyses were conducted because we did not find that FC estimates from the Yeo17 atlas explained any variance in the psychological metrics, and we thought that this might be because the Yeo17 atlas has large parcels. Notably, prior work has shown that more fine-grained atlases provide greater predictive utility than more coarse-grained ones [64]. The results of this analysis demonstrate that even with FC estimates using a more fine-grained atlas, FC does not explain a notable amount of variance in the psychological metrics under study in the current work.

- Correlations between FC estimates from the Schaefer 400 atlas and each of the psychological metrics were estimated. None of the comparisons survived FDR correction. This analysis was conducted to supplement the analysis detailed above.
